# FreeSurfer version-shuffling can boost brain age predictions

**DOI:** 10.1101/2024.06.14.599070

**Authors:** Max Korbmacher, Lars T. Westlye, Ivan I. Maximov

## Abstract

- The influence of FreeSurfer version-dependent variability in reconstructed cortical features on brain age predictions is average small when varying training and test splits from the same data.
- FreeSurfer version differences can lead to some variability in brain age dependent on the choice of algorithm and individual differences in brain morphometry, highlighting the advantage of repeated random train-test splitting.
- Shuffling of differently processed FreeSurfer data dependent on the FreeSurfer version increases performance and generalizability of the brain age prediction model.

## Background

The concept of brain age is based on the prediction of participants’ age from a set of brain images using machine learning algorithms.^1–4^ Subtracting brain age from a person’s chronological age, also called brain age gap or brain age delta, has been outlined as a promising biomarker of brain health, where a larger gap has been related to poorer health outcomes for the participant.^2,4^ While brain age estimation has been widely applied, there are still multiple sources of prediction errors, which cannot simply be attributed to biological brain features, such as in-scanner subject motion^5^, field strength^6^ or other MRI-related technical features, or machine learning workflows (feature space, algorithm, bias correction choices)^7–10^.

Software packages used in image processing such as FreeSurfer^11^ present version-dependent variability in image derived estimates, such as cortical thickness in T_1_-weighted metrics.^12–15^ The naturally arising question is then whether and how this source of variability influences brain age predictions.

A more pronounced knowledge of brain age variability induced by MRI processing software is key for inference and further comparability of brain age study results, as well as the application of trained models on unseen data processed with different FreeSurfer versions. Hence, we aimed to explore brain age predictions based on data obtained from FreeSurfer versions 5.3.0 and 7.4.1^a^ using the latest data release of the UK Biobank (UKB). Moreover, we introduce version-shuffling for brain age predictions, leveraging training and test data from different FreeSurfer versions boosting prediction accuracy and robustness.

## Methods

### Participants

We included N = 4,395 UKB participants free of neurological and psychiatric disorders (ICD-10 categories F, G, I, and stroke), and removed MRI feature-level outliers defined as 5 standard deviations above and below the mean. The participants were scanned at three sites in the UK: Cheadle (55.38%), Newcastle (34.99%), and Reading (9.62%). The participants’ age ranged from 49.33 to 82.84 years of age (Mean±SD=64.84±7.27), including 52.53% females.

### Magentic Resonance Imaging

T_1_-weighted images were obtained at 3T on a Siemens Skyra 3T running VD13A SP4 (as of October 2015), with a standard Siemens 32-channel radio frequency receive head coil using a 3D MPRAGE sequence (sagittal, in-plane acceleration iPAT=2, prescan-normalise) at 1x1x1 mm^3^, field-of-view: 208x256x256 matrix, taking approximately 5 minutes.

### Feature derivation

Cortical reconstruction was performed using FreeSurfer versions 5.3.0 and 7.4.1 (from here on v5 and v7), extracting regional cortical thickness, volume and surface area using the Desikan-Killiany cortical parcellation scheme on a single T_1_-weighted 3D image per participant.^16^ The data were processed on the Sigma2 Norwegian national computation cluster on RedHat Linux Release 8.10 (Ootpa), Linux kernel version: 4.18.0-513.24.1.el8_9.x86_64.

### Power analysis

With the conservative assumption of a large parameter shrinkage of 40%, we estimated a required training sample size of N = 2,009 to train a brain age model on the selected M = 204 grey matter features using the pmsampsize package^17^ in R, version 4.3.3. Hence, we could be confident that our sample size was appropriate to train a meaningful model when using a 50-50 split of the sample, to firstly train models for each FreeSurfer version using least absolute shrinkage and selection operator (Lasso)^17^, Support Vector Machine (SVM)^19^ regression, eXtreme Gradient Boosting (XGBoost)^20^ regression, and Light Gradient Boosting Machine (LightGBM)^21^, and conventional linear regression.

### Main statistical analyses

We then predicted the age in the test set within and between FreeSurfer version-specific models (see Fig.1 for the procedure). For completeness, in addition to Lasso, we applied alternative machine learning algorithms, namely, SVM^19^ regression, XGBoost^20^ regression, and LightGBM^21^ regression, as well as linear regression. For comparability, we kept all hyperparameters constant at default settings, except for Lasso, where we implemented a higher maximum number of iterations (i_Lasso_= 100,000) to reach optimal alpha tolerance. A single train-test iteration, from here on the *initial iteration*, served to compare model performance. For simplicity, the following analyses are focused on results from the best performing model only.

**Fig.1:**
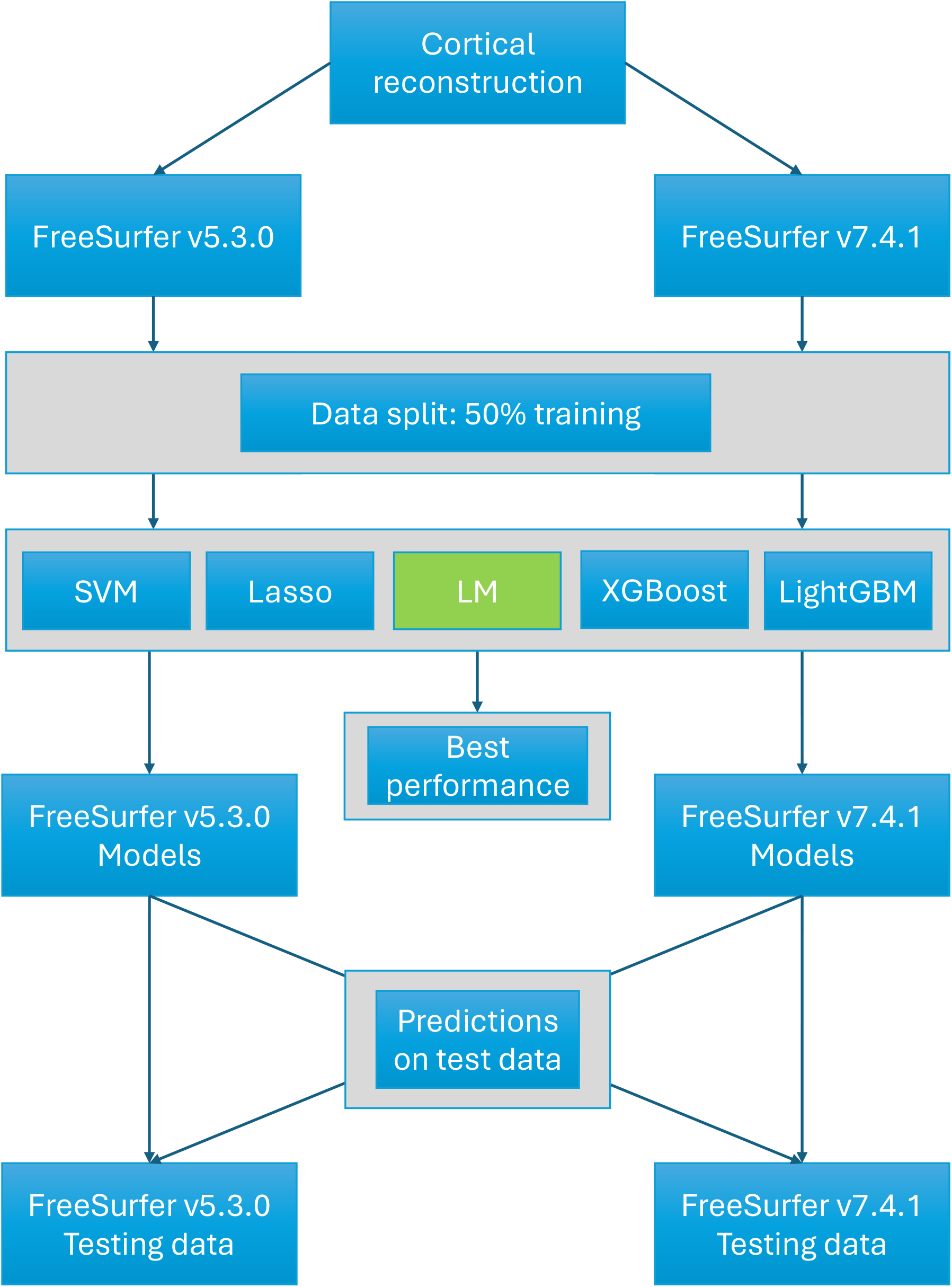
Data processing scheme. Acronyms of the used ML algorithms: SVM = Support Vector Machine, Lasso = Least absolute shrinkage and selection operator, LM = Linear Model (Linear Regression), XGBoost = eXtreme Gradient Boosting, LightGBM = Light Gradient Boosting Machine.

To account for individual differences, we used i = 1,000 permutations to randomly split training from test data into equal parts (50% each). Then, we investigated models trained on version-shuffled data, i.e. model training based on features of both FS5 and FS7 in the *initial iteration*, which each training and test sets were composed of 50% FS5 and 50% FS7 data. As a final step, we varied both train-test splits and version specific feature composition in i = 1,000 permutations, where the data was again equally split into train and test data, and each of these sub-sets contained 50% FS5 and 50% FS7 features.

We examined common error metrics (root mean squared error (RMSE), mean absolute error (MAE)), the coefficient of determination (R^2^), the Akaike information criterion (AIC), the Bayesian information criterion (BIC), and the Pearson correlations between chronological and predicted brain age. Moreover, we estimated differences between these correlations, using Zou^22^ and Hittner tests^23^, as well as the version-dependent marginal means of the predicted ages accounting for sex, age, and scanner site, using simple linear models. We also examined scanner site closer as a potential covariate of interest when using different models to predict on the same data using linear mixed effects models where the version of the training data was the random intercept, sex and age covariates, and site the fixed effect. To give a practical example of how training and test data can influence study outcomes, we estimated marginal sex differences from linear models predicting brain age from age, sex, and site for each combination of training and test sets.

### Analyses of the data structure

Supplemental analyses aided to further explore potential reasons for the observed variability, we examined a) the correlations between FS5 and FS7 features, b) the correlations of the most important features, ranked by permutation feature importance, between versions, c) age correlations at the feature level, d) executed a Principal Component Analysis (PCA) sorting the features into thickness, area, volume.

## Results

### Cross-version predictions

In the *initial iteration*, we evaluated the performance of different machine learning algorithms against each other. SVM, XGB, LightGBM, and Lasso yielded similar results as the linear regression models, yet with lower prediction performance (see Supplement 1-2). In other words, linear regression was outlined as the superior model. For illustration, we present the predictions of this first iteration in Fig.2. Note that the differences between the predictions were statistically significant^b^.

**Fig.2:**
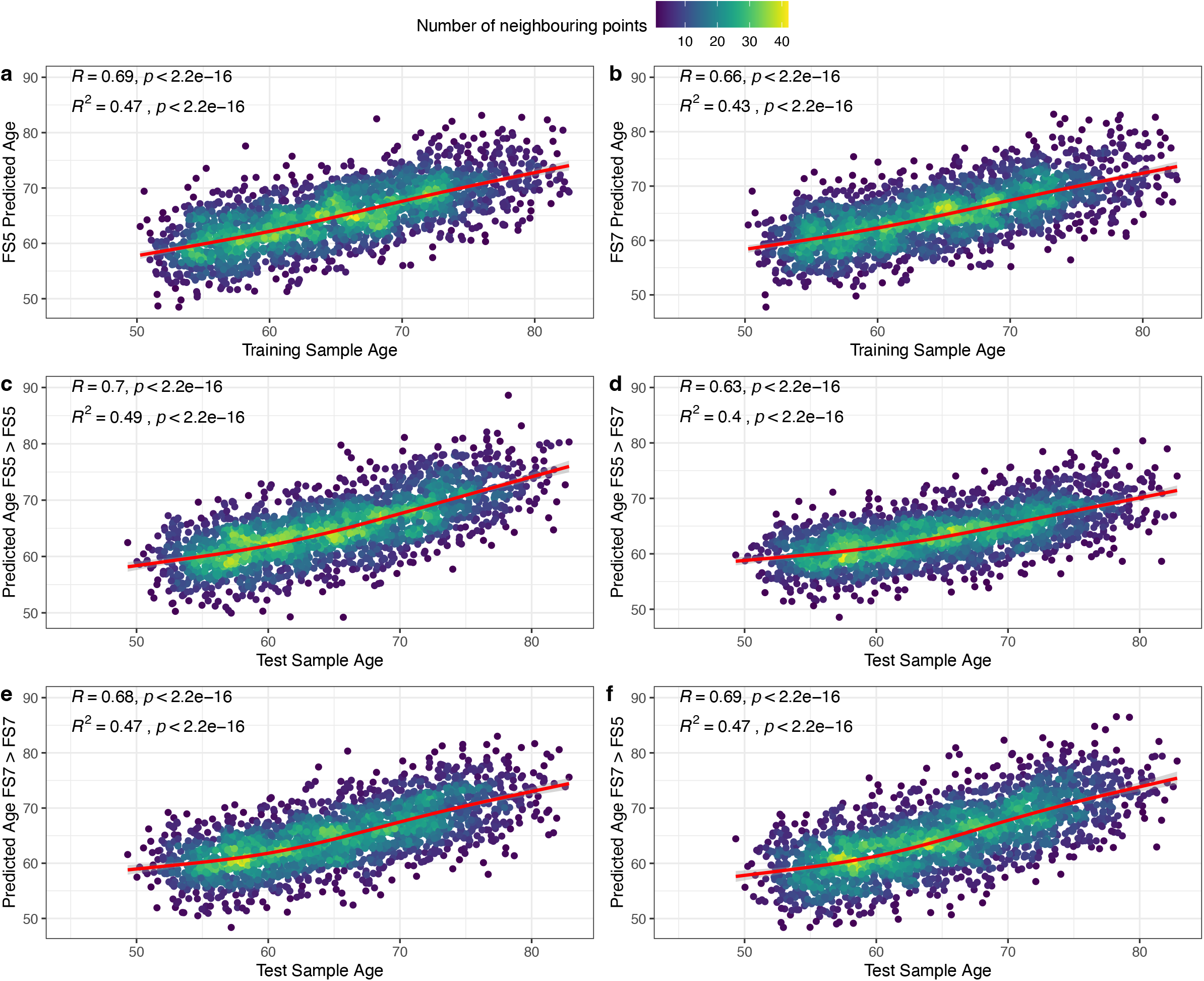
Training and test performance for linear regression models trained on different FreeSurfer versions in the *initial iteration*. Panels a-b: prediction performance on training data. Panels c-f: prediction performance on test data. FS5 = FreeSurfer v5, FS7 = FreeSurfer v7, arrows indicate which trained model was used to predict which training data, e.g.: “FS7 > FS5” indicates that a model trained on FreeSurfer v7 processed features was used to predict age derived by FreeSurfer v5 processed features. The red line shows a smooth cubic fit with k = 4 knots.

When randomly sampling training and test data, we obtained an acceptable model performance in both training and test data (r > 0.67, R^2^ > 0.39, MAE < 4.51 years, RMSE < 9.03 years, see Table 1), comparable to brain age studies using a similar set of FreeSurfer features^c^. The variability in the findings from the i = 1,000 random train-test splits suggest that there is an influence of individual differences in brain morphometry on brain age predictions from FS5 and FS7 (Table 1).

**Table 1:**
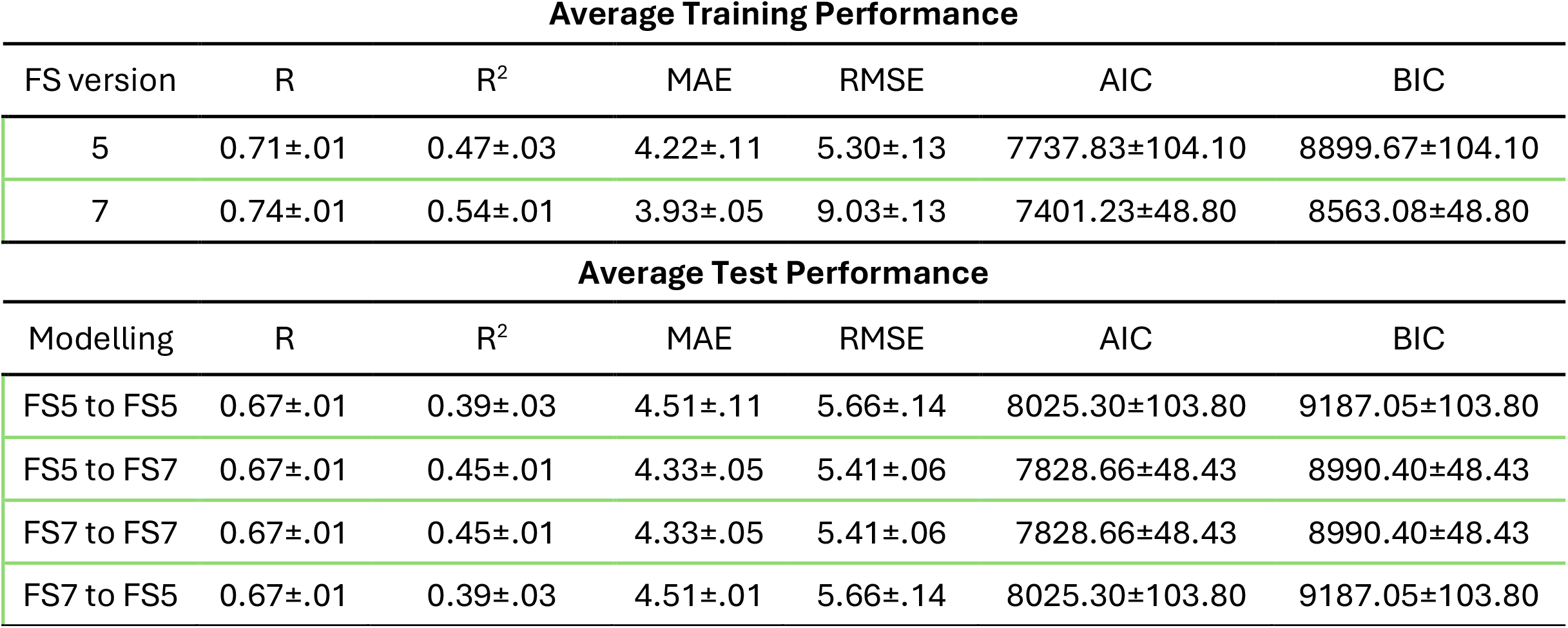
Version-specific linear regression train-test performance across i = 1,000 iterations for random training and test splitting. R = Pearson’s Correlation coefficient, adjusted R^2^ = adjusted variance explained (linear model), adjusted R^2^ corrected = adjusted variance explained (linear model correcting for sex and scanner site), MAE = mean absolute error, RMSE = root mean squared error, AIC = Akaike information criterion, BIC = Bayesian information criterion. FS = FreeSurfer. RMSE and MAE are indicated in years. Values are presented as Mean ± Standard Deviation. *Note*: The composition of the 50-50 train-test splits were varied (i = 1,000) to examine the influence of individual differences. Training-test performance of the second-best performing algorithm, Lasso, can be found in Supplement 5.

**Average Training Performance**
| FS version | R | R<sup>2</sup> | MAE | RMSE | AIC | BIC |
| --- | --- | --- | --- | --- | --- | --- |
| 5 | 0.71±.01 | 0.47±.03 | 4.22±.11 | 5.30±.13 | 7737.83±104.10 | 8899.67±104.10 |
| 7 | 0.74±.01 | 0.54±.01 | 3.93±.05 | 9.03±.13 | 7401.23±48.80 | 8563.08±48.80 |

**Table 1:** Version-specific linear regression train-test performance across $i = 1,000$ iterations for random training and test splitting.
| Modelling | R | R<sup>2</sup> | MAE | RMSE | AIC | BIC |
| --- | --- | --- | --- | --- | --- | --- |
| FS5 to FS5 | 0.67±.01 | 0.39±.03 | 4.51±.11 | 5.66±.14 | 8025.30±103.80 | 9187.05±103.80 |
| FS5 to FS7 | 0.67±.01 | 0.45±.01 | 4.33±.05 | 5.41±.06 | 7828.66±48.43 | 8990.40±48.43 |
| FS7 to FS7 | 0.67±.01 | 0.45±.01 | 4.33±.05 | 5.41±.06 | 7828.66±48.43 | 8990.40±48.43 |
| FS7 to FS5 | 0.67±.01 | 0.39±.03 | 4.51±.01 | 5.66±.14 | 8025.30±103.80 | 9187.05±103.80 |

### Version-shu?ling

Version-shuffled models performed better than single-version models on version-shuffled data, and on cross-version predictions (Table 2). Version-shuffled models performed also comparable to single-version models predicting on the data from the *initial iteration* of the same version as the single-version models (Table 2) and provided slightly better predictions in general in comparison to single-version models where the train test split composition was varied (compare Tables 1-2), also when varying both version-specific feature composition and train-test composition (compare Table 1 and Supplement 6).

**Table 2:** Lasso Modelling Performance Across FreeSurfer Versions averaged across the i = 1,000 trained models. R = Pearson’s Correlation coefficient, adjusted R^2^ = adjusted variance explained (linear model), adjusted R^2^ corrected = adjusted variance explained (linear model correcting for sex and scanner site), MAE = mean absolute error, RMSE = root mean squared error. FS = FreeSurfer. RMSE and MAE are indicated in years. Modelling column: Mix = version-shuffled, FS5 = FreeSurfer v5 only, and FS7 = Freesurfer v7 only. Values are presented as Mean ± Standard Deviation. 1 Refers to randomly assembled version-shuffled training data. 2 Refers to randomly assembled version-shuffled test data. 3 Refers to models which were established on a single iteration (*i* =1), predicting on constant data from the *initial iteration*. *Note:* The 50-50 train-test split was held constant but features from FS5 and FS7 were varied within the splits (50-50 FS5 and FS7) to probe the influence of version-shuffling. Hence, there was only a single iteration i = 1, being the initial iteration for single version predictions, as reflected in Fig.2 (as also noted in footnote 3 in this figure).

| Modelling | R | R<sup>2</sup> | MAE | RMSE | AIC | BIC |
| --- | --- | --- | --- | --- | --- | --- |
| Mix to Mix<sub>train</sub><sup>1</sup> | 0.73±.003 | 0.54±.004 | 3.98±.02 | 4.97±.02 | 7433.21±20.65 | 8594.30±20.65 |
| Mix to Mix<sub>test</sub><sup>2</sup> | 0.68±.004 | 0.46±.006 | 4.25±.03 | 5.31±.03 | 7770.85±26.54 | 8933.34±26.54 |
| FS5 to Mix<sub>test</sub><sup>2</sup> | 0.66±.004 | 0.42±.007 | 4.43±.03 | 5.50±.03 | 7929.20±24.78 | 9091.69±24.78 |
| FS7 to Mix<sub>test</sub><sup>2</sup> | 0.68±.003 | 0.45±.005 | 4.28±.02 | 5.36±.02 | 7816.03±20.38 | 8978.52±20.38 |
| Mix<sup>3</sup> to FS5<sub>test</sub> | 0.70±.003 | 0.48±.004 | 4.17±.02 | 5.24±.02 | 7710.38±18.87 | 8872.87±18.87 |
| FS5 to FS5<sub>test</sub> | 0.70 | 0.49 | 4.17 | 5.20 | 7511.61 | 8218.22 |
| FS7 to FS5<sub>test</sub> | 0.69 | 0.44 | 4.30 | 5.41 | 7691.80 | 8398.42 |
| Mix<sup>3</sup> to FS7<sub>test</sub> | 0.67±.003 | 0.45±.005 | 4.32±.02 | 5.38±.02 | 7831.58±18.01 | 8994.07±18.01 |
| FS5 to FS7<sub>test</sub> | 0.63 | 0.36 | 4.69 | 5.80 | 8001.73 | 8708.34 |
| FS7 to FS7<sub>test</sub> | 0.68 | 0.46 | 4.25 | 5.32 | 7621.26 | 8327.87 |

Zou^22^ and Hittner tests^23^ comparing correlations between brain age and age from the initial iteration for FS5 and FS7 models (i = 1) and the average across predictions (i.e., bagging) when varying the composition of the version-shuffled data (as in Table 2) showed no differences between within single-version predictions (FS5 to FS5 and FS7 to FS7) and version-shuffled models’ predictions (Table 3). However, version-shuffled models outperformed single-version models on all other accounts.

**Table 3:** Comparison of correlations between age and predicted age in single- and version-shuffled predictions. Version-shuffled predictions were averaged across the i = 1,000 predictions to account for variations in version-dependent feature compositions. For comparability, single version estimates are from the initial iteration (i = 1) to hold the training-test sample structure constant. rdi’ refers to the difference in correlations between age and predicted age. CIlower and CIupper refer to the Zou’s^22^ 95% confidence interval constructed around rdi’. Z and p values indicate Hittner test^23^ derived Z-statistic and according (uncorrected) p-value. The modelling pairs indicate which correlations between each predicted age and chronological age were compared.

| Modelling Pairs | $r_{diff}$ | $CI_{lower}$ | $CI_{upper}$ | Z | p | $p_{FDR}$ |
| --- | --- | --- | --- | --- | --- | --- |
| Mix to FS5 & FS5 to FS5 | 0.004 | -0.002 | 0.010 | 1.412 | 0.158 | 0.172 |
| Mix to FS5 & FS7 to FS5 | 0.018 | 0.012 | 0.024 | 6.214 | $5.17 \times 10^{-10}$ | $2.07 \times 10^{-9}$ |
| FS5 to FS5 & FS7 to FS5 | 0.014 | 0.003 | 0.025 | 2.549 | 0.011 | 0.019 |
| Mix to FS7 & FS5 to FS7 | 0.042 | 0.034 | 0.051 | 10.281 | $< 2.22 \times 10^{-16}$ | $< 2.22 \times 10^{-16}$ |
| Mix to FS7 & FS7 to FS7 | -0.007 | -0.012 | -0.001 | -2.298 | 0.022 | 0.0323 |
| FS5 to FS7 & FS7 to FS7 | -0.049 | -0.062 | -0.036 | -7.458 | $8.82 \times 10^{-14}$ | $5.29 \times 10^{-13}$ |
| Mix to Mix & FS5 to Mix | 0.017 | 0.010 | 0.023 | 5.329 | $9.89 \times 10^{-8}$ | $1.98 \times 10^{-7}$ |
| Mix to Mix & FS7 to Mix | 0.006 | 0.001 | 0.011 | 2.240 | 0.025 | 0.033 |
| FS5 to Mix & FS7 to Mix | -0.011 | -0.021 | 0.000 | -1.965 | 0.049 | 0.059 |
| Mix to Mix & FS5 to FS5 | 0.004 | -0.003 | 0.011 | 1.048 | 0.295 | 0.295 |
| Mix to Mix & FS7 to FS7 | 0.023 | 0.016 | 0.031 | 6.071 | $1.27 \times 10^{-9}$ | $3.05 \times 10^{-9}$ |
| FS5 to FS5 & FS7 to FS7 | 0.023 | 0.016 | 0.031 | 6.071 | $1.27 \times 10^{-9}$ | $3.05 \times 10^{-9}$ |

The observed differences in correlations between chronological age and brain ages can also be observed in the same pattern as presented in Table 3 when directly comparing marginal means of brain ages in version pairs using simple linear models correcting for age, sex and site (Table 4). We observe version-dependent brain age differences of up to 1.42 years using models trained on different versions to predict on the same data.

**Table 4:**
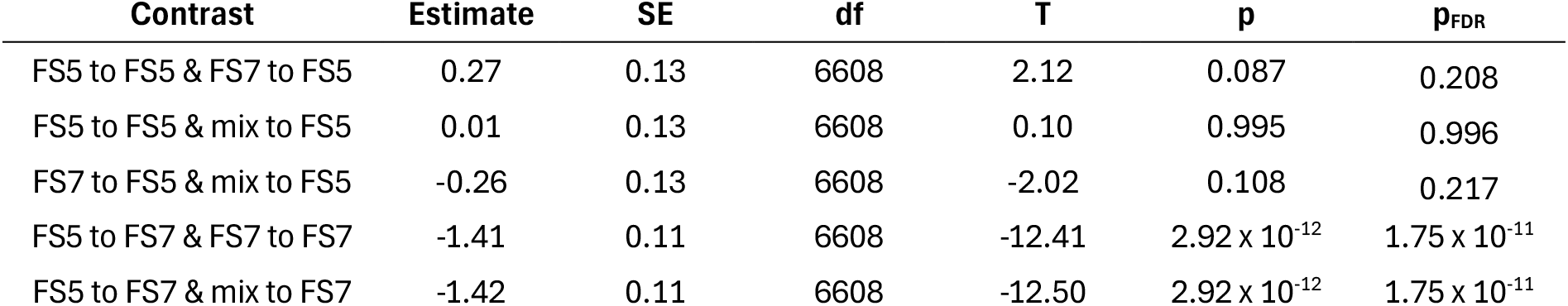
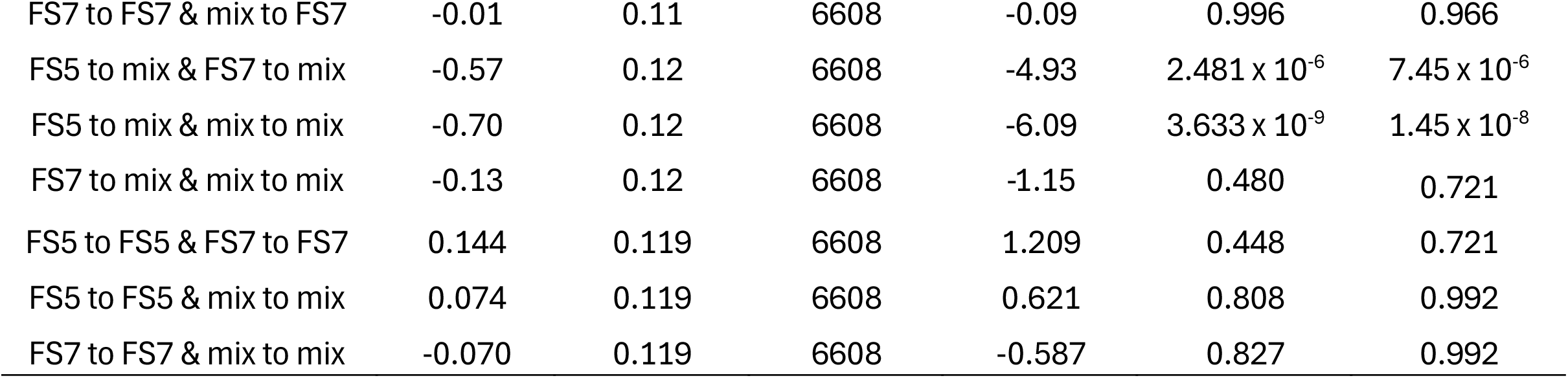
Marginal mean comparison of different models’ brain age predictions corrected for age, sex and site. Estimate = difference between brain ages (in years). SE = Standard Error (in years). df = degrees of freedom. Raw and false discovery corrected p-values are indicated. Version-shuffled brain age predictions were averaged across the i = 1,000 predictions to account for variations in version-dependent feature compositions. For comparability, within and between single version brain age predictions were used from the initial iteration (i = 1) to hold the training-test sample structure constant. For a test of the effect of *both* train-test splits *and* (version-dependent) feature composition see Supplement 6.

Linear mixed effect models, with the FreeSurfer version being the random intercept, and scanner site, age and sex being fixed effects, did not suggest an effect of scanner site on brain age estimates (p > .340).

To provide a practical example of how the different training and test data compositions can affect study results, such as group differences, we used simple linear models for each of the nine brain age predictions (representing brain age models trained on either FS5, FS7 and version-shuffled data predicting into the respective data). The results suggest partly opposing estimates of group differences (Figure 2).

### Analyses of the data structure

a) Correlations between the same features estimated by the different FreeSurfer versions were on average Pearson’s correlation coefficients’ Mean r_train_ = 0.88 in the training data and Mean r_test_ = 0.88 in the test data. To examine potential confounding effects of age, we additionally estimated the same version correlations for participants grouped by age decade, resulting in consistent correlations across age-bins (Supplement 4). The smallest correlations (r < 0.69) were observed for enthorinal cortex as well as temporal and frontal poles’ volume and surface area. b) The 10 highest ranked features in the permutation feature importance rankings of models trained on FS5 or FS7 data were similar for the models trained on FS5 compared to FS7 (Supplement 3). Moreover, these features were strongly correlated between versions (r > 0.90). The strongest age-associated *regions* in the training data reflected prominent features contributing to the age predictions (inferior parietal, superior frontal, and precentral) when ranked by permutation feature importance (see Supplement 3), in terms of thickness across versions (r < -0.42), but smaller for volume (r < -0.22), and small for surface area (r < - 0.07). c) Moreover, feature-age associations were on average influenced by the Freesurfer versions, as indicated by paired samples t-tests (Mean Pearson’s r: [FS5_train_ = -0.17, FS7_train_ = -0.15, t = -4.998, p = 1.25 x 10^-6^, d = -0.350], [FS5_test_ = -0.18, FS7_test_ = -0.13, t = -14.859, p < 2.20 x 10^-16^, d = -1.040]), yet d) PCAs produces similarly structured components across FreeSurfer versions (see Figure 4 for an overview of a-d).

**Fig.3.**
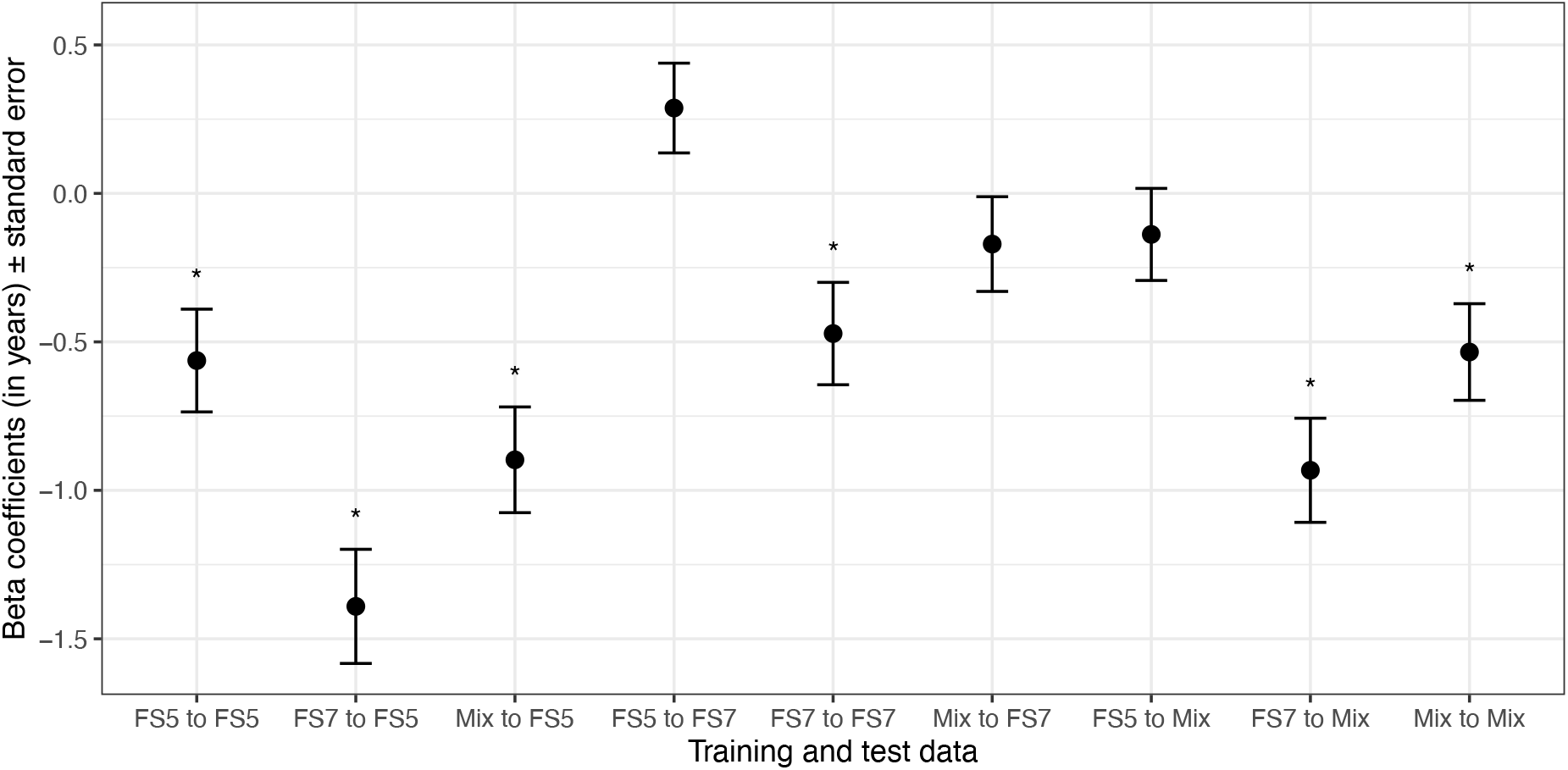
Age and site-corrected sex differences in brain age estimates from models trained on and predicting on data from FS5, FS7, and a version mix. Coefficients b > 0 indicate a higher brain age of females, b < 0 for males, respectively. The sex differences were significant at pFDR < .05 for all predictions except for FS5 to FS7, mix to FS7, and FS5 to mix. FS5 = FreeSurfer version 5, FS7 = FreeSurfer version 7, Mix = 50-50 mix of FS5 and FS7 data. FDR = false discovery rate. Version-shuffled models were established by randomly sampling training data in *i* = 1,000 iterations, and version-shuffled test data were established in the same procedure. The average of the predictions (from *i* > 1) was used here (bagging approach).

**Fig.4:**
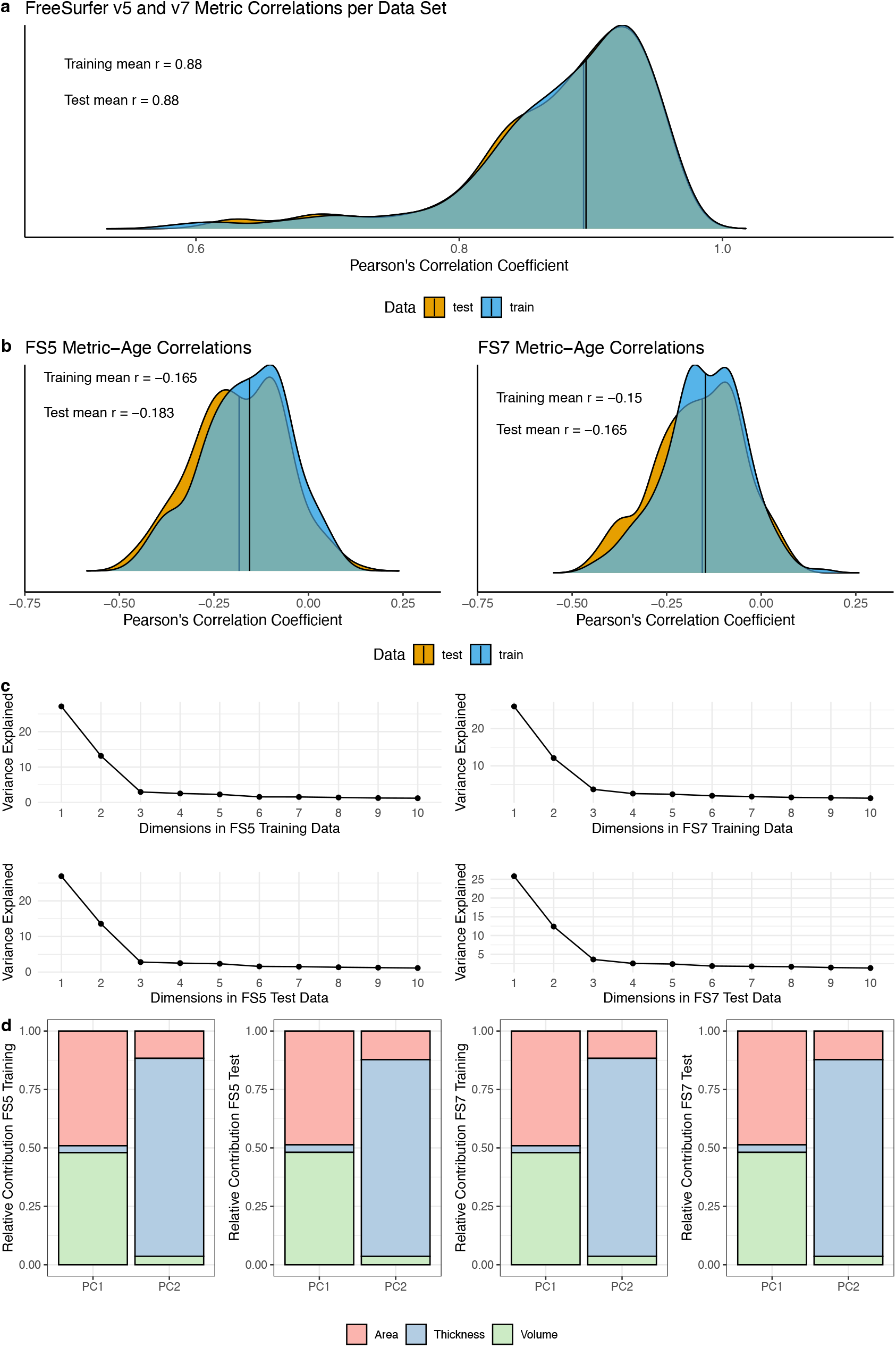
Panel a) Shows the correlations of each feature estimated from FS5 and FS7. Panel b) presents age associations of each feature by data set (and FreeSurfer version). Panel c) indicates the variance explained for each of the first 10 components for each dataset. Panel d) presents the relative distributions of feature types for the first two components.

## Discussion

Brain age predictions are influenced by various factors, some of which are not biological. For meaningful predictions and construction of brain age biomarkers, such factors need to be mapped and potentially controlled for. Here, we examined how different FreeSurfer versions might influence brain age predictions, comparing versions 5.3.0 and 7.4.1. While it is known that FreeSurfer version differences lead to variability in feature estimates, these differences seem to be relatively small. As brain age is a multivariate method, such systematic differences might however influence model training and predictions.

While we find that the differences between brain age predictions across FreeSurfer versions are negligible when using i = 1,000 iterations to randomly split training and test sets (Table 1), in individual cases these predictions can induce significant performance differences (here described on the example of one individual *initial iteration*; Fig.2-3; Table 2-4). This can be both expressed in terms of direct differences in brain age estimates (of up to 1.42 years; Table 2-4), as well as resulting non-negligible differences in estimations of the effect of sex (Figure 3).

We present that training on features from different versions might boost model performance and generalizability. Brain age is predicted most accurately across examined models when training on version-shuffled data and predicting on version-shuffled data (Table 1-2). Moreover, predictions from models trained on version-shuffled data on single-version data show only small differences to prediction from models trained on that same single version (Table 4). Specifically, version-shuffled models’ age predictions are just as good as single-version models when predicting on the specific single version data considering FS5 (FS5_train_ to FS5_test_) and nearly as good as single-version models considering FS7 (FS7_train_ to FS7_test_). Version-shuffled models clearly outperform single-version models attempting to predict age from data processed with another FreeSurfer version (FS5_train_ to FS7_test_ or FS7_train_ to FS5_test_; Table 3). Moreover, version-shuffled models outperform single-version models in predicting age in version-shuffled data, with this difference being however only marginally significant for predictions from the FS7-trained model^d^. Finally, even when varying test and training splits and which features were recruited from which FreeSurfer version (Supplement 6), version-shuffled models performed similar to, yet slightly better than, single-version models (Table 2-3), outlining the advantages of version-shuffling when attempting to build generalizable models.

These findings are important considering goals such as increasing accuracy, generalizability, and at the same time the continuous updating of FreeSurfer leading to variability of versions in the data available to researchers. Particularly when attempting to train brain age models, which is usually done on large and multi-site datasets, often requiring merging data processed in different software version, computational overhead can be reduced by shuffling FreeSurfer versions during the training process (when different versions are used at random) instead of reconstructing cortical features from scratch to match a certain target version. Such advantage of reduced version-specific feature engineering is also true when considering the usage of pre-trained models with the goal of predicting brain age in new samples.

The outlined version-shuffling procedure is additionally rewarded by the largest accuracy across models when predictions are made on version-shuffled test data (Table 2), especially when using a bagging approach where predictions from multiple models are averaged (Table 3-4).

Moreover, we find simple linear regression to outperform more advanced machine learning approaches in this ageing sample, older than 49 years of age. This might speak to previously outlined near-linear relationships between age and brain features in that age group^1,3^, in contrast to non-linear relationships considering the whole lifespan^26,27^. Linear models have a simple structure, are fast to compute, and might be more generalizable across FreeSurfer versions than other algorithms such as Lasso, which was the second-best performing algorithm (compare Table 1 and Supplement 5).

To provide additional information on differences in data structures resulting from the usage of FS5 compared to FS7, we ran supplementary analyses comparing feature estimates across versions, their age-associations, as well as principal components. The principal components/correlational structure of features was similar across versions, and cross-version feature correlations were strong (r > 0.87). The lowest correlations between the same features estimated from different FreeSurfer versions were located in the frontal and temporal poles, which is coherent with previous findings comparing similar or different FreeSurfer versions.^12,15^ Additionally, feature-age associations differed between versions, potentially also influencing brain age predictions. In particular, age-associations of FS5 features were stronger than for FS7, also shown previously by Haddad and colleagues^15^.

While brain age estimates from FS5 features were stronger associated with age than predictions based on FS7 features in the initial iteration (i = 1; Figure 2), this was not the case when repeatedly randomly splitting the data into training and test sets, showing no differences in predictions on test data (Table 1). FS7 models from that first iteration produced better cross-version and version-shuffled predictions than FS5. Hence, FS7 might be the better choice for accurate brain age predictions across versions or on version-shuffled data. A few explanations for the version-dependent variability can be found in version-dependent differences in feature estimations (and their age-dependencies), which finally lead to differences in feature contributions to the age predictions (Supplement 3). However, such feature rankings are limited by multicollinearity in the present data and must be interpreted with care. Nevertheless, the outlined differences in both predictions (Figure 1, Table 2-4) and accompanying feature importances (Supplement 3) in the *initial iteration* compared to an absence of prediction performance differences when varying training and test splits (Table 1) suggest an influence of individual differences on FreeSurfer version-specific cortical reconstructions. Repeated random train-test splits, as executed here, might hence produce more robust brain age estimates. Another, and potentially preferable, option is to use version-shuffling, where the composition of version-specific features can be varied, if data from more than one version are available for the same participants. Both strategies are computationally feasible when either using a) a fixed selection of hyperparameters for more complex models, such as Lasso, SVM, or tree boosting, or b) simple models such as linear regression. This allows for predictions from hundreds or thousands of models which can for example be averaged to provide more robust brain age estimates (bagging procedure).

The attempt to establish brain age as a biomarker entails to establish robust associations of brain age with pheno- and genotypes or pheno- and genotype-based group differences. Here, we used sex differences as an example phenotype. Sex-differences in brain age usually entail a higher brain age in males in samples with comparable age spans^1,28^, which largely corresponds with our findings (Figure 3). However, we find that when predicting on FS7 data with an FS5-trained model, this relationship can even be reversed. Yet, considering the estimated derivatives, within-version sex differences (FS5_train_ to FS5_test_, FS7_train_ to FS7_test_, FSmix_train_ to FSmix_test_) were highly similar. In contrast, sex differences in cross-version predicted brain ages (FS5_train_ to FS7_test_ or FS7_train_ to FS5_test_) deviated from within-version estimates more strongly than when obtained from version-mixed models (FSmix_train_ to FS7_test_ or FSmix_train_ to FS5_test_). These sex differences in brain ages indicated by models trained on version-shuffled data were laying somewhat between the estimates from single-version trained models. This indicates that cross-version predictions might lead to over- or underestimations of pheno- or genotype effects on brain age. Yet, there is no clear answer to which model supplies most accurate sex differences. The reason is that there is no ground truth for these differences. The goal of these analyses was also not to benchmark sex differences in brain age. Instead, with the presented sex differences, we wanted to highlight that choices of the FreeSurfer version when computing training and testing data features can impact phenotype associations with brain age or group differences.

There are various limitations to this study. First, we assumed an equal share of FS5 to FS7 features within both training and test sets. Different compositions of features of different versions are more realistic and can be further examined. Second, site effects are a known phenomenon, influencing both FreeSurfer-derived metrics^15^ and potentially interaction effects of version and site on brain age estimation. While we did not find site effects, both site effects and site-version interactions require further investigations. These analyses could focus on a wider range of versions and especially consider data sets with larger heterogeneity between sites, for example in terms of acquisition protocols. Third, the manuscript is limited to the Desikan-Killiany atlas and its containing regions. Other atlases and features derived from other procedures, such as vertex-wise analyses, should be examined in future investigations. Connected to this point, there should be, fourth, observed different samples beyond the UK Biobank as well as additional phenotypes. Fifth, much of the inference presented here to compare brain age models trained on and predicting on version-shuffled and single-version data is based on a single random iteration of train-test splits (besides the findings presented in Table 1). This choice was based on the computational overhead which would result from varying training-test splits and their feature compositions (e.g., 1,000 x 1,000 times) as well as non-trivial analyses strategies resulting from the generated data. However, we included an analysis where we varied both train-test splits and feature compositions together in i = 1,000 iterations (Supplement 6). Sixth, in an optimal scenario, features would be available for more than one version for the entire sample to being able to randomly draw training and test splits assembled of the data from the different FreeSurfer versions. In reality, most likely not all data are available for several versions, and instead systematically per version. For example, one site processed all data with FS5, another site with FS7, or all data before a certain date were processed with an earlier FreeSurfer version than at a later point in time. We did not examine such systematic version influences, but rather at random and with equal contributions.

FS7-trained models outperform FS5-trained models predicting on version-shuffled data, FS7 data, and differ only slightly when predicting on FS5 data. This suggest FS7 as the preferrable version when choosing single-version training data. However, models trained on version-shuffled data perform just as good FS5 trained models when predicting on FS5 data, differ only slightly from FS7-trained models when predicting on FS7 data, and are clearly outperforming on single-version trained models when predicting on version-shuffled data. Models trained on version-shuffled data might hence be more generalizable considering variability in versions and might pose an advantage over single-version trained models.

## Code Availability

All code is available at https://github.com/MaxKorbmacher/FS_BrainAge

## Data Availability

All raw data are available from the UKB (https://www.ukbiobank.ac.uk) in accordance with data acessing procedures.

## Ethics Statement

This study was conducted using UKB data under Application 27412. UKB received approval from the North West Multi-centre Research Ethics Committee (MREC) as a Research Tissue Bank.

## Acknowledgements

We want to thank the facilitators and all participants of the UKB.

This research was funded by the Research Council of Norway (#223273, L.T.W.); and the European Union’s Horizon2020 Research and Innovation Programme (#802998 L.T.W.). The work was performed on the Service for Sensitive Data (TSD) platform, owned by the University of Oslo, operated and developed by the TSD service group at the University of Oslo IT-Department (USIT). Computations were performed using resources provided by UNINETT Sigma2—the National Infrastructure for High Performance Computing and Data Storage in Norway.

## Author Contributions

M.K.: Conceptualisation, Data curation, Formal Analysis, Methodology, Project Administration, Software, Visualization, Writing – original draft, Writing – review & editing.

L.T.W: Funding acquisition, Writing – review & editing.

I.I.M.: Conceptualisation, Data curation, Funding acquisition, Software, Writing – review & editing

## Footnotes

a Multiple changes have been made from Freesurfer version 5.3.0 to 7.4.1 to improve both efficiency and accuracy of the reconstructions, such as changes in denoising, white-grey-matter boundary identification, and surface reconstruction.

b While the prediction performance was on average similar between FreeSurfer versions, we observed differences in predictions in the practical example of the *initial iteration* (Fig.2). Here, Zou^22^ and Hittner tests^23^, used to compare correlations between brain age and age, indicated that FS5 predicted more accurately within version, in training data (r_diff_ = 0.069, 95%CI[0.049, 0.085], z = 7.272, p < .0001), and test data (r_diff_ = 0.019, 95%CI[0.006, 0.033], z = 4.493, p < .001), and FS7 between versions (r_diff_ = -0.054, 95%CI[-0.072, -0.036], z = -5.9637, p < .0001; see also Fig.2, Supplement 1-2).

c Examples: Beck et al.^24^: R^2^ = 0.73, MAE = 7.2 years, RMSE = 9.11 years; Han et al.^25^: R^2^ = 0.47, MAE = 7.50.

d Using a bagging procedure averaging across predictions from different models.

## Supplement

**Supplement 1:** Training Model Performance Across Algorithms (*initial iteration*)

| R | adjusted R<sup>2</sup> | adjusted R<sup>2</sup> (corrected) | MAE | RMSE | FreeSurfer version | Model |
| --- | --- | --- | --- | --- | --- | --- |
| 0.56 | 0.31 | 0.32 | 4.90 | 6.11 | 5 | XGBoost |
| 0.38 | 0.15 | 0.21 | 5.60 | 6.76 | 5 | SVM |
| 0.60 | 0.36 | 0.36 | 4.80 | 5.90 | 5 | LightGBM |
| 0.65 | 0.42 | 0.42 | 4.52 | 5.58 | 5 | Lasso |
| **0.69** | **0.47** | **0.47** | **4.27** | **5.33** | **5** | **LM** |
| 0.53 | 0.28 | 0.28 | 5.07 | 6.27 | 7 | XGBoost |
| 0.33 | 0.11 | 0.17 | 5.76 | 6.92 | 7 | SVM |
| 0.57 | 0.32 | 0.33 | 4.98 | 6.06 | 7 | LightGBM |
| 0.62 | 0.39 | 0.39 | 4.63 | 5.73 | 7 | Lasso |
| **0.66** | **0.43** | **0.43** | **4.41** | **5.52** | **7** | **LM** |

Modelling Performance Across FreeSurfer Versions and Algorithms in Training Data. R = Pearson's Correlation coefficient, adjusted R<sup>2</sup> = adjusted variance explained (linear model), adjusted R<sup>2</sup> corrected = adjusted variance explained (linear model correcting for sex and scanner site), MAE = mean absolute error, RMSE = root mean squared error. RMSE and MAE are indicated in years.

**Supplement 2:**
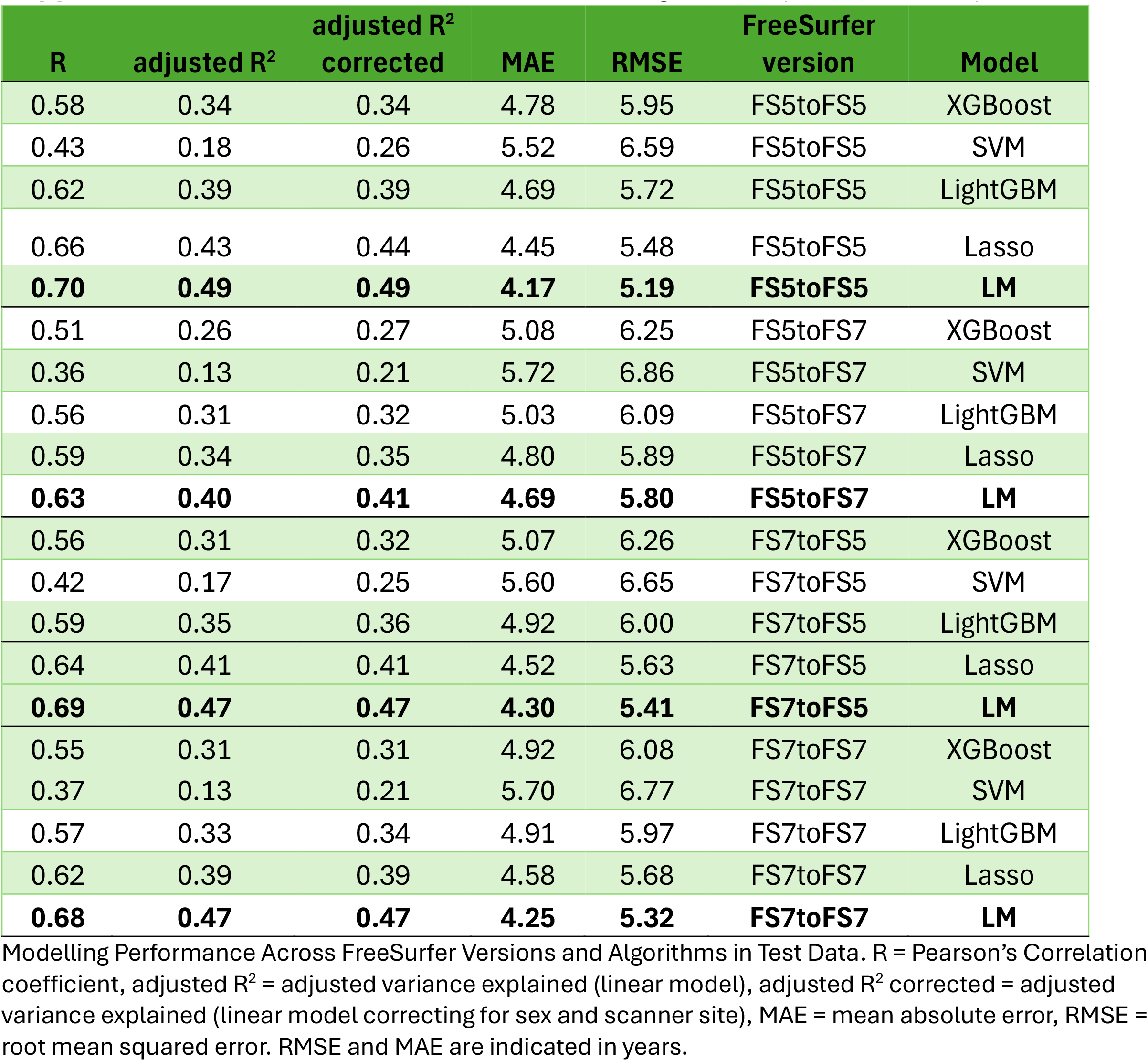
Test Model Performance Across Algorithms (*initial iteration*)

**Supplement 3:** Permutation feature importance: top 10 features and their variance explained when predicting age (*initial iteration*)

| FreeSurfer v5 | FreeSurfer v7 |
| --- | --- |
| lh_superiorfrontal_area 0.341867 +/- 0.007639 | lh_precentral_volume 0.233111 +/- 0.008399 |
| lh_inferiorparietal_area 0.275286 +/- 0.011307 | lh_precentral_area 0.210369 +/- 0.007713 |
| lh_inferiorparietal_volume 0.260479 +/- 0.009025 | rh_posteriorcingulate_area 0.194168 +/- 0.006988 |
| rh_inferiortemporal_area 0.253444 +/- 0.010406 | lh_bankssts_volume 0.165247 +/- 0.005948 |
| rh_inferiortemporal_volume 0.243288 +/- 0.007102 | lh_bankssts_area 0.162048 +/- 0.010068 |
| lh_superiorfrontal_thickness 0.209069 +/- 0.010630 | lh_medialorbitofrontal_area 0.157978 +/- 0.012232 |
| lh_supramarginal_volume 0.207518 +/- 0.005090 | lh_medialorbitofrontal_volume 0.153799 +/- 0.007422 |
| lh_supramarginal_area 0.183548 +/- 0.008399 | rh_inferiortemporal_area 0.148451 +/- 0.006095 |
| lh_precentral_volume 0.181147 +/- 0.006269 | lh_superiorfrontal_area 0.133032 +/- 0.005665 |
| lh_precentral_area 0.151394 +/- 0.006774 | rh_cuneus_volume 0.120757 +/- 0.005527 |

**Supplement 4:**
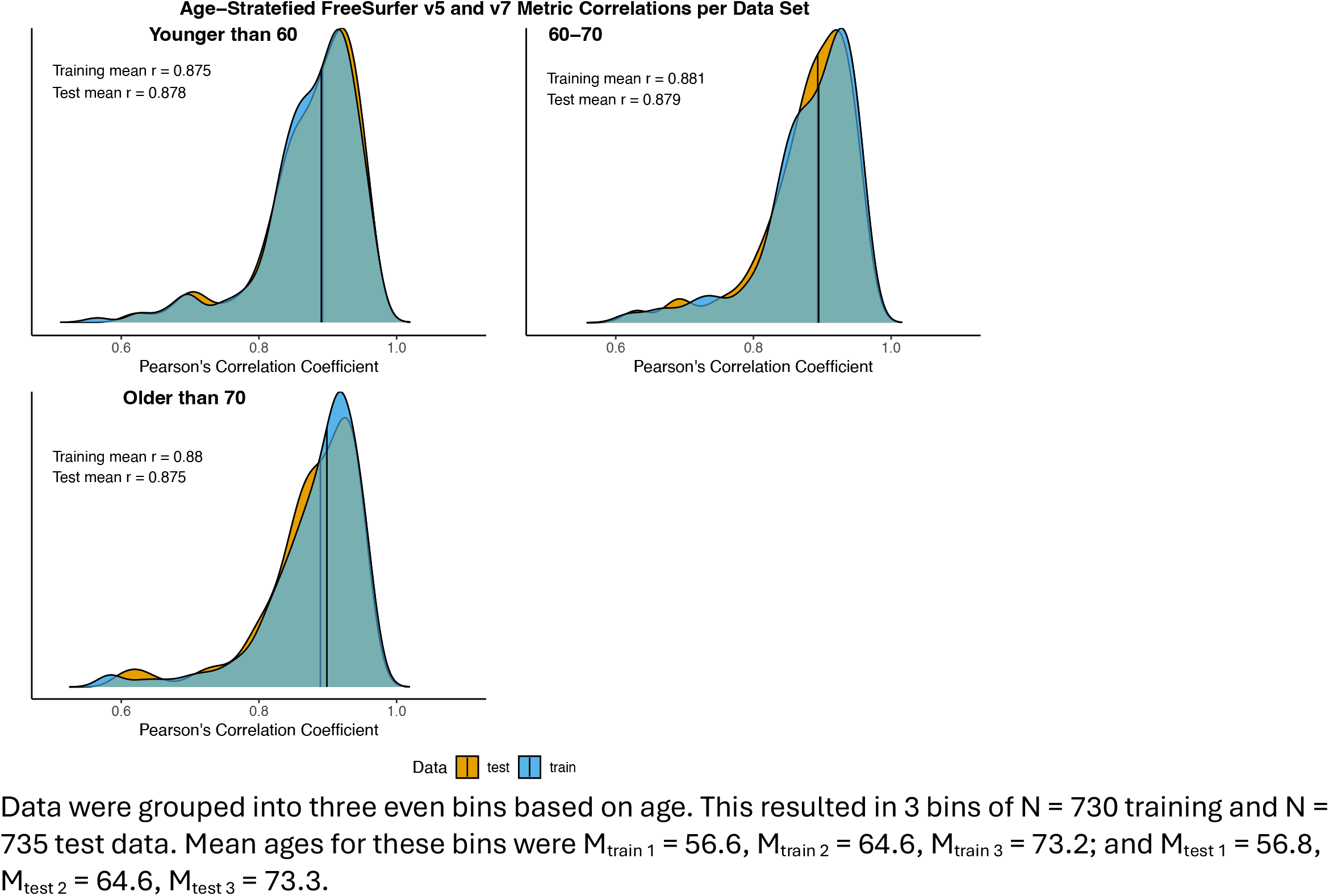
Age stratefied correlations between FS5 and FS7 feature estimates.

**Supplement 5:** Lasso training and test when 1000 times repeatedly randomly sampling training and test data (compare with Table 1)

| Average Training Performance |  |  |  |  |
| --- | --- | --- | --- | --- |
| R | R<sup>2</sup> | MAE | RMSE | FS version |
| 0.67±.01 | 0.43±.01 | 4.41±.06 | 5.47±.07 | 5 |
| 0.67±.01 | 0.45±.01 | 4.34±.05 | 5.37±.06 | 7 |
| Average Test Performance |  |  |  |  |
| R | R<sup>2</sup> | MAE | RMSE | Modelling |
| 0.63±.01 | 0.38±.02 | 4.62±.07 | 5.73±.08 | FS5 to FS5 |
| 0.62±.01 | 0.38±.01 | 4.62±.05 | 5.72±.06 | FS5 to FS7 |
| 0.62±.01 | 0.38±.01 | 4.62±.05 | 5.72±.06 | FS7 to FS7 |
| 0.62±.01 | 0.38±.02 | 4.62±.07 | 5.73±.08 | FS7 to FS5 |

**Supplement 6:** Supplement 6: Performance of models varying both training and test splits as well as feature composition (FreeSurfer version)

|  | **R2** | **r** | **MAE** | **RMSE** | **AIC** | **BIC** |
| --- | --- | --- | --- | --- | --- | --- |
| Mean train | 0.548 | 0.740 | 3.919 | 4.890 | 7384.931 | 8546.773 |
| SD train | 0.010 | 0.007 | 0.050 | 0.058 | 52.120 | 52.120 |
| Mean test | 0.449 | 0.674 | 4.324 | 5.395 | 7813.901 | 8975.650 |
| SD test | 0.013 | 0.008 | 0.054 | 0.063 | 51.580 | 51.580 |

We used i = 1,000 iterations to randomly split the data into equal parts of training and test data and each of these portions into equal parts of FS5 and FS7 features for training and testing.

